# Functional cerebellar connectomes interfacing motor adaptation and reinforcement feedback

**DOI:** 10.64898/2026.03.25.714278

**Authors:** Martina Bracco, Candice Appriou, Vridhi Rohira, Xavier Corominas-Teruel, Alice Person, Odirachukwunma Orah, Francois-Xavier Lejeune, Salim Ouarab, Benoît Béranger, Karim N’Diaye, Yulia Worbe, Traian Popa, Antoni Valero-Cabre, Cecile Gallea

**Affiliations:** Mov’it Team, Sorbonne Université, Institut du Cerveau - Paris Brain Institute - ICM, Inserm, CNRS, APHP, Hôpital de la Pitié Salpêtrière, 47 Bd de l’Hôpital, Paris, 75013, France; Dynamics, Plasticity and Rehabilitation Group, Frontlab Team, Sorbonne Université, Institut du Cerveau - Paris Brain Institute - ICM, Inserm, CNRS, APHP, Hôpital de la Pitié Salpêtrière, 47 Bd de l’Hôpital, Paris, 75013, France; Danish Research Centre for Magnetic Resonance, Centre for Functional and Diagnostic Imaging and Research, Copenhagen University Hospital Hvidovre, Kettegård Alle 36, 2650, Hvidovre, Denmark; Data Analysis Core (RRID: SCR_026138), Sorbonne Université, Institut du Cerveau - Paris Brain Institute - ICM, CNRS, APHP, Hôpital de la Pitié Salpêtrière, 47 Bd de l’Hôpital, Paris, 75013, France; Center for NeuroImaging Research (CENIR), Sorbonne Université, Institut du Cerveau - Paris Brain Institute - ICM, Inserm, CNRS, APHP, Hôpital de la Pitié Salpêtrière, 47 Bd de l’Hôpital, Paris, 75013, France; PRISME, Sorbonne Université, Institut du Cerveau - Paris Brain Institute - ICM, Inserm, CNRS, APHP, Hôpital de la Pitié Salpêtrière, 47 Bd de l’Hôpital, Paris, 75013, France; MySpace Lab and NeuroRehab Research Center, Service of University Neurorehabilitation (SUN), Lausanne University Hospital, Institution of Lavigny and University of Lausanne, 5 Av. Pierre-Decker, 1011 Lausanne, Switzerland; Lake Lucerne Institute (LLUI), 9 Rubistrasse, Vitznau, 6354, Switzerland

**Author notes:** These authors contributed equally to this work (Antoni Valero-Cabre, Cecile Gallea).

**Keywords:** Cerebellum, Motor Adaptation, Reinforcement Learning, Functional Connectivity, Dopamine, Serotonin

## Abstract

Motor adaptation is driven by sensory prediction errors, yet reinforcement feedback can alter the speed and retention of adaptive behaviors. The cerebellum is central in motor adaptation, but posterior lobules, especially Lobule VI (CB6) and Crus I (CBcrus1), also participate in reinforcement-related signaling. Serotoninergic and dopaminergic systems, key modulators of motivational processes, directly influence cerebellar activity. However, how these neuromodulatory systems contribute to cerebellar network organization and to reinforcement-based adaptation in humans remains unclear. We employed a multimodal framework (Receptor-Enriched Analysis of functional Connectivity by Targets; REACT) using PET-derived Serotonin/Dopamine Transporter (SERT/DAT) templates to enrich resting-state functional Magnetic Resonance Imaging (rs-fMRI). We estimated SERT- and DAT-enriched functional connectivity from CB6 and CBcrus1 within motor-adaptation and reinforcement-learning networks, and tested their correlations with performance in a visuomotor adaptation task completed under reward and punishment. Our findings show lobule-specific neuromodulatory organization within the cerebellum. CB6 exhibited predominantly DAT- and SERT-enriched connectivity with motor adaptation networks, while CBcrus1 showed stronger SERT-enriched communication extending to both motor and reinforcement networks. Crucially, distinct cerebello-cortical neuromodulatory networks predicted individual differences in adaptation rate. DAT-enriched CBcrus1 connectivity was linked to punishment-driven adaptation, whereas SERT-enriched cerebello-orbitofrontal connectivity predicted faster adaptation across both reward and punishment contingencies. Furthermore, we observed overlaps between SERT- and DAT-enriched networks in the medial orbitofrontal area for Crus I, which predicted retention following punishment, underscoring the role of convergent neuromodulation in stabilizing adapted movements. We conclude that partially segregated yet convergent cerebello-cortical networks support interactions between motor and motivational behaviors, with combined and opposing effects of serotonergic and dopaminergic neuromodulation accounting for the speed and retention of adaptive behaviors.

## INTRODUCTION

Voluntary actions are fundamental for achieving goals in diverse and often unpredictable environments. To maintain goal pursuit amidst this complexity, the brain must continuously adapt upcoming actions to minimize error. This process, known as motor adaptation, supports movement recalibration based on sensory feedback and the individual’s current state to ensure flexible, effective performance [Bastian, 2008]. Motor adaptation is classically driven by sensory prediction errors; however, growing evidence suggests that this process is also modulated by reinforcement feedback. When motor adaptation is associated with feedback of positive or negative value (reward or punishment), both the rate of adaptation and the retention of learned responses are affected [Galea et al., 2015; Mongeon et al., 2013; Quattrocchi et al., 2017; Venkatakrishnan et al., 2011]. For instance, previous work has shown that adaptation is faster following punishment, but the adapted movements are retained longer after reward [Galea et al., 2015]. Yet, the brain mechanisms that support the interaction between adaptation and reinforcement feedback remain unclear.

The cerebellum has long been recognized as the central structure for motor adaptation [Miall and Wolpert, 1996; Shadmehr and Krakauer, 2008; Taylor and Ivry, 2014]. Cerebellar lobules participate in multiple motor and non-motor processes and are embedded within distinct large-scale cortical networks [Bracco et al., 2025; Diedrichsen et al., 2019; D’Mello et al., 2020; Nettekoven et al., 2024]. Crucially, functional heterogeneity also exists within individual lobules. For example, lobule VI (CB6), which plays a prominent role in motor adaptation [Hardwick et al., 2013; Johnson et al., 2019], exhibits classic motor features such as somatotopic organization, but also engages cognitive functions such as working memory and attention [Diedrichsen et al., 2019]. In contrast, cerebellar Crus1 (CBcrus1) is more strongly coupled to associative cortices and is preferentially engaged in higher-order cognitive processes, consistent with a rostro–caudal gradient in cerebellar functional organization [Buckner et al., 2011; Diedrichsen et al., 2019; D’Mello et al., 2020]. Despite these functional differences, both regions have been implicated in reinforcement feedback processing [Chen et al., 2022; Kostadinov and Häusser, 2022; Kruithof et al., 2023; Manto et al., 2024; O’Doherty et al., 2003; Tobler et al., 2005; Wagner et al., 2017]. This functional overlap raises a critical question: do cerebellar networks contribute to the interaction between motor adaptation and reinforcement feedback through shared functional representations, or through distinct but interacting neural pathways?

Reinforcement learning is shaped by interacting neuromodulatory systems. Reward and punishment engage distinct yet overlapping neural circuits [Garrison et al., 2013; Monosov and Hikosaka, 2012; Oldham et al., 2018; Wächter et al., 2009], shaped by the single and interactive contributions of dopamine and serotonin. Dopamine is strongly associated with reward-related learning, whereas serotonin has been implicated in both reward and, more prominently, punishment processing [Cools et al., 2008; Cools et al., 2009; DenOuden et al., 2013; Doira et al., 2004; Palminteri et al., 2009; Palminteri et al., 2012; Worbe et al., 2011]. Critically, recent work in rats has shown that coordinated interaction between striatal dopamine and serotonin neurons is crucial for promoting associative learning and long-term retention [Cardozo Pinto et al., 2025]. These findings indicate that dopamine and serotonin can exert both complementary and distinct influences within the same system.

Both serotonergic and dopaminergic systems directly modulate cerebellar activity. The cerebellum receives serotonergic input from reticular and raphe nuclei [Bishop and Ho, 1985; Dieudonné, 2001; Pierce et al., 1977; Schweighofer et al., 2004]. This input modulates neuronal firing rates and synaptic plasticity, essential for learning [Fleming and Hull, 2018; Lippiello et al., 2016; Di Matteo et al., 2008; Murano et al., 2011]. While the contribution of dopamine to cerebellar activity is more controversial [Oertel, 1993], positron emission tomography (PET) studies using dopamine transporter (DAT) ligands have shown substantial levels of dopamine in the primate cerebellum [Glaser et al., 2006; Varrone et al., 2009], particularly in CBcrus1 and dentate nucleus ([Flace et al., 2021]; for review). Immunohistochemical studies further report the presence of dopaminergic receptors in the cerebellum [Khan et al., 1998; Khan et al., 2000; Melchitzky and Lewis, 2000], and its role in the neuromodulation of cerebellar motor and non-motor functions [Cutando et al., 2022; Locke et al., 2018; Yu et al., 2009]. In addition, recent resting-state functional magnetic resonance imaging (rs-fMRI) mapping of monoaminergic nuclei further suggests that the cerebellum is a shared large-scale connectivity target of both serotonergic (e.g., dorsal raphe nucleus) and dopaminergic (ventral tegmental area – VTA, and substantia nigra pars compacta – SNc) systems, consistent with broader overlap across monoaminergic functional networks [Saiz-Masvidal et al., 2025]. Together, these findings position these neurochemical systems to influence cerebellar communication based on sensory prediction errors and reinforcement feedback [Kilteni and Ehrsson, 2024; Kilteni and Henrik Ehrsson, 2020; Kruithof et al., 2023; Parr et al., 2021].

In the present study, we investigated the joint and distinct contributions of serotonergic and dopaminergic systems within different cerebellar networks to reinforcement-based motor adaptation. Investigating these systems in humans is challenging, as they require separate, costly, and invasive PET procedures for each neurotransmitter system. To overcome these constraints, we employed a multimodal approach integrating high-resolution PET-derived templates of DAT [García-Gómez et al., 2013] and the serotonin transporter (SERT; [Beliveau et al., 2017]) with rs-fMRI analyses [Dipasquale et al., 2019]. This approach yields neurotransmitter-informed functional networks enabling the examination of how large-scale connectivity patterns are modulated by serotonin and dopamine, without directly measuring neurotransmitter release or signaling.

We specifically focused on CB6 and CBcrus1 as regions of interest (ROIs), given their established roles in motor adaptation and reinforcement-related processing [Chen et al., 2022; Johnson et al., 2019]. Because these cerebellar regions exhibit distinct functional connectomes [Diedrichsen et al., 2019; Nettekoven et al., 2024; O’Reilly et al., 2009], we hypothesized that they would show partially segregated SERT- and DAT-enriched functional networks (Hypothesis 1). At the same time, these networks could converge in cortical regions where sensorimotor and prefrontal systems interact, consistent with potential integration between motor and reinforcement-related processes (Hypothesis 2). Finally, we hypothesized that individual performance in reinforcement-based motor adaptation would be differentially related to SERT- and DAT-enriched cerebellar networks. Adaptation rate under punishment and reward would preferentially associate with separable cerebellar nodes and neurotransmitter systems, whereas retention would reflect more integrated serotonergic and dopaminergic contributions (Hypotheses 3A–B).

## MATERIAL AND METHODS

### Population

We recruited 24 right-handed (as defined by the Edinburgh Questionnaire; [Oldfield, 1971]) healthy volunteers (13 females, 25.8 ± 5.9 years; mean ± SD). All participants reported no history of medical, neurological, or psychiatric conditions, had normal or corrected-to-normal vision, and met the additional MRI safety criteria [Shellock and Spinazzi, 2008]. Written informed consent was obtained from all participants. The study was approved by the Île de France I ethics committee (CPPIDF1-2021-ND92-cat.2) and conducted in accordance with the Declaration of Helsinki.

### Experimental design and data acquisition

Participants first underwent an rs-fMRI session, followed by two behavioral sessions in which they performed a reinforcement-based motor adaptation task. The behavioral sessions were identical, with the exception that the task was performed under a different reinforcement contingency (reward or punishment). To prevent savings from prior sessions from affecting subsequent performance, behavioral sessions were separated by at least 10 days. The order of contingencies was randomized across participants to control for order effects.

#### Resting-state fMRI

MRI data were acquired on a 3T Siemens Prisma scanner (Siemens Healthineers, Erlangen, Germany) using a 64-channel head coil. A mirror was attached to the head coil so that participants could see a display presented in front of them. Participants were instructed to lie still with their eyes open, fixating on a cross presented at the center of the display, and to remain awake and relaxed without performing any specific task. Head motion was minimized using foam cushions.

For each participant, we recorded a T1-weighted MP2RAGE with the following parameters: repetition time (TR) = 5000 ms, echo time (TE) = 3.24 ms, inversion times TI1 = 700 ms and TI2 = 2500 ms, flip angles FA1 = 4° and FA2 = 5°, field of view (FOV) = 256 × 256 mm², matrix = 256 × 256, 176 sagittal slices, slice thickness = 1 mm, resulting in an isotropic voxel size of 1 × 1 × 1 mm³.

Resting state functional data were collected using a multi-echo multi-band T2*-weighted gradient-echo echo-planar imaging (EPI) sequence with the following parameters: TR = 1660 ms, TEs = 14.2 ms / 35.39ms / 56.58ms, flip angle = 74°, FOV = 210 × 210 mm², matrix = 84 × 84, 60 axial slices, slice thickness = 2.5 mm, no gap, voxel size = 2.5 × 2.5 × 2.5 mm³, in-plane acceleration GRAPPA=2, and a multiband acceleration factor of 3. One run lasted 12 minutes (440 volumes). The acquisition was repeated twice with a 20-minute break between runs to increase data quality and avoid participant drowsiness. Single-band reference and inverted phase-encoding reference images were acquired for distortion correction and accurate registration.

#### Reinforcement-based motor adaptation task

Participants performed a joystick-based reaching task with their dominant right hand (Logitech Attack 3, Logitech Europe S.A., Lausanne, Switzerland). Stimuli were displayed on a computer screen (grey background, viewing distance of 800 mm), controlled via MATLAB (The MathWorks, Natick, MA, USA) and Psychophysics Toolbox. Movements were recorded at 140 Hz. Participants had to shoot through a target using a cursor controlled by the joystick.

Each trial began with the appearance of a white fixation cross (20 mm height/length) at the center of the screen, superimposed on a larger outer circle (open white, 320 mm diameter) also centered on the screen. A white cursor (16 mm diameter) then appeared at the central position, indicated by an open white circle (20 mm diameter). This was followed by the presentation of a target (open white circle, 20 mm diameter) at one of four possible locations along the upper half of the outer circle (20°, 60°, 120°, 160°). Targets were presented pseudo-randomly so that each set of four consecutive trials (a cycle) contained one instance of each target position.

Participants were instructed to control the cursor with the joystick and perform fast, shooting movements to hit the target. When the cursor crossed the outer circle, it disappeared, and the intersection point was marked with a yellow circle (16 mm diameter) to denote the endpoint, while the target turned green (hit) or red (miss). This prompted participants to passively return the joystick to the central location. Movements shorter than 100 ms or longer than 600 ms triggered on-screen feedback (“Too fast” / “Too slow”). The next trial began after a 4000 ms interval (jittered between 3500–4500 ms), and only after the participant had maintained the cursor within the central circle for at least 200 ms.

After familiarization, each session consisted of eight blocks of 48 trials each (divided into 12 cycles). The session began with two baseline blocks, one with continuous visual cursor feedback and one without cursor feedback (only the start and final positions were visible). In the third to the fifth block, the cursor trajectory was rotated relative to the actual joystick trajectory (*Adaptation*), requiring participants to adjust their reaching movements to hit the target. Rotation directions (either 25° clockwise (+25°) or 30° counterclockwise (−30°)) were identical within a session but were alternated between sessions and randomized across participants. Finally, in blocks six to eight, the rotation was removed, and participants performed three blocks where the cursor was hidden (as in the second baseline block) to assess post-effects of the adaptive behavior (*Retention*).

During the three adaptation blocks, participants received reinforcement feedback contingent on performance (reward or punishment). In the reward condition, accurate performance was rewarded by earning points, while in the punishment condition, inaccurate performance resulted in loss of points. Accuracy was defined as the angular distance between the cursor and target position, and reward and punishment were implemented as follows:

**Reward:** participants started with 0 points, and better performance gained more points (+4 for a hit, +3 for <10° error, +2 for <20° error, +1 for <30° error, 0 for ≥30° error)

**Punishment:** participants started with the maximum number of points (456), and better performance decreased point loss (0 for a hit, −1 for <10° error, −2 for <20° error, −3 for <30° error, −4 for ≥30° error)

Participants received trial-by-trial feedback and cumulative scores at the end of each block.

### Data analysis

#### Resting-state fMRI

##### Preprocessing

Structural (T1-weighted) images were first denoised using the MP2RAGE background correction algorithm ([O’Brien et al., 2014]; https://github.com/benoitberanger/mp2rage), then segmented and normalized to the Montreal Neurological Institute (MNI) space using the Computational Anatomy Toolbox (CAT12). Resting-state echo-planar imaging (EPI) images were first preprocessed with AFNI [Cox, 1996] for slice-timing correction (reference slice = first slice, t = 0) using a 7th-order Lagrange polynomial interpolation, and motion correction using rigid body transformations (no susceptibility correction; reference scan fixed with the MIN_OUTLIER option in afni_proc.py). Despiking was then performed using the AFNI 3dDespike function. The correction of individual distortions was performed using the inverted phase-encoding reference images with the TOPUP function of FSL. Subsequently, TEDANA combined signal across echoes and used a principal component analysis to separate blood-oxygen-level dependent BOLD from non-BOLD components based on the echo time dependence of the BOLD component3 (non-BOLD components were visually inspected for each individual). Denoised rs-fMRI images were then co-registered to individual structural scans using rigid-body transformation in SPM12, and normalized to MNI space via mutual information cost function and 4th-degree B-spline interpolation. The normalization to the MNI space was applied to the structural image and then to the rs-fMRI image. Finally, data were spatially smoothed (FWHM = 4 mm) and temporally band-pass filtered (0.01–0.1 Hz) using the standard procedure implemented in SPM12.

##### Regions and networks of interest

Individual cerebellar lobules were segmented from each participant’s anatomical T1-weighted image using the Spatially Unbiased Infratentorial Template (SUIT) toolbox (version 3.7; [Diedrichsen, 2006; Diedrichsen et al., 2009]). The resulting cerebellar maps were normalized to MNI space, and CB6 and CBcrus1 were extracted as seed ROIs. To select the target networks, we considered the regions identified by meta-analyses for motor adaptation [Johnson et al., 2019] and reinforcement learning [Chen et al., 2022]. The motor adaptation network included bilateral frontal eye field, middle frontal gyrus, supplemental motor area (SMA), precentral gyrus (primary motor and premotor areas), postcentral gyrus, superior parietal lobule, inferior parietal lobule, parietal operculum and posterior insula. The reinforcement learning network contained the bilateral medial orbitofrontal cortices (MOFC), anterior cingulate cortices (ACC), nucleus accumbens (NAC), and VTA.

#### Neurotransmitter-Enriched Network Analysis

We applied the REACT, a framework that integrates PET-derived maps of receptor/transporter density with resting-state fMRI to estimate neurotransmitter-enriched functional connectivity [Dipasquale et al., 2019]. High-resolution in vivo atlases of the 5-HTT SERT and DAT were used to enrich rs-fMRI analyses with the spatial distribution of these proteins in the healthy brain. These atlases were created from molecular and structural high-resolution neuroimaging data consisting of PET and MRI scans acquired in 210 and 30 healthy individuals, respectively [Beliveau et al., 2017; García-Gómez et al., 2013]. As we focused on cerebellar networks, both templates were normalized by scaling the map values between 0 and 1, considering the gray matter of the cerebellum segmented with SUIT. Because the cerebellum is often used as the reference region for kinetic modeling of SERT, absolute values in this region are typically lower than those in cortical or striatal areas. Nevertheless, measurable variability in SERT density exists across cerebellar lobules [Beliveau et al., 2017]. To retain this relative variability, normalization was performed separately for SERT and DAT by scaling each map to the maximum value within the cerebellum.

The REACT procedure then involved a two-step regression analysis using the general linear model (GLM) [Dipasquale et al., 2019]. In the first step, we used the cerebellar SERT and DAT maps as spatial regressors to estimate neurotransmitter-enriched functional connectivity (FC) by fitting BOLD fluctuations across voxels to the spatial distribution of each map [Griffanti et al., 2015]. These regressions were implemented in FSL (fsl_glm) using a region-of-interest approach restricted to CB6 and CBcrus1. To match the spatial resolution of the receptor maps and preserve the spatial information of the cerebellar receptor, rs-fMRI images were upsampled to 1 × 1 × 1 mm3. In the second step, the participant-specific time series were used as temporal regressors in a second regression to estimate the participant-specific whole-brain maps of the BOLD response. At this stage, we used the rs-fMRI images sampled at 2 x 2 x 2 mm3. The analysis was conducted on the whole grey matter volume for CB6 and CBcrus1 separately, using the GLM in SPM12 with SERT and DA maps as regressors of interest, defining a minimum of 26 nuisance regressors: six motion parameters and their first, second and third derivatives, as well as white matter and CSF signals extracted with the TAPAS toolbox [Frässle et al., 2021].

### Statistical Analysis

For each participant, we computed the positive effect of each regressor of interest (SERT and DAT) to generate participant-specific spatial maps. These maps were then entered into a second-level analysis to examine the specificity of cerebellar functional networks. We used a flexible factorial design in SPM12 with two within-subject factors (*Lobule*: CB6, CBcrus1; *Receptor Type*: SERT, DAT) with repeated measures. F contrasts tested the main effects of *Lobule* and *Receptor Type*, and the *Lobule* x *Receptor Type* interaction, with post-hoc t-tests evaluating the direction of significant effects. In particular, comparisons between CB6 and CBcrus1 tested whether their SERT-and DAT-enriched network distributions were spatially segregated (Hypothesis 1). Additionally, we conducted a conjunction analysis of SERT- and DAT-enriched maps within each lobule to identify overlapping cortical and subcortical regions modulated by both receptor types (Hypothesis 2). Statistical significance was set at *p* < 0.05 with Family-Wise Error (FWE) correction applied at the level of the network (adaptation, reinforcement; see “Regions and network of interest” section).

### Behavioral data

#### Preprocessing

We quantified participants’ performance as reach angle, defined as the angle of the hand’s movement relative to the target position (0°; MATLAB; The MathWorks, Natick, MA, USA). During the adaptation blocks, a visuomotor rotation (e.g., −30° or +25°) was applied, requiring participants to produce a compensatory reach angle (e.g., +30° or −25°), i.e., higher values indicate better performance. To make performance comparable across both rotation directions, we used the absolute deviation from the required angle. During the retention blocks where no rotation was applied, the goal was to produce a 0° reach angle, i.e., lower values indicate better performance. Trials were excluded if the reach time was too fast (<100 ms) or too slow (>600 ms), or if the initial reach angle deviated by more than 60° from the target. Valid trials were averaged into consecutive blocks of four trials (cycle); missing points were linearly interpolated.

#### State-Space model analysis

To characterize behavioral performance, we fitted state-space models [Galea et al., 2015; Smith et al., 2006] separately to the adaptation and retention phases for each participant and feedback contingency (reward, punishment).

##### Adaptation Model (AB model)

This model assumed that the current motor state, denoted as 𝑧_𝑛_, is updated based on two components: the previous motor state (𝑧_𝑛−1_) and the performance error experienced in the previous cycle (𝑟_𝑛_ − 𝑧_𝑛−1_), with denoting the trial number. The model was defined as:

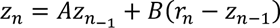

Here, *A* represented the retention factor, capturing the proportion of the previous motor state that is retained from one cycle to the next. *B* was the adaptation rate, indicating how strongly the system adjusts in response to the error on the previous cycle.

##### Retention Model (AC Model)

This model assumed that retention occurred as an exponential decay of the motor state with a persistent after adaptation. It was defined as:

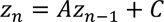

Here, *B* as fixed to zero, meaning that no adaptive process occurred because the cursor was hidden and no trajectory feedback was available. *C* represented a fixed bias that persisted over cycles, independent of recent performance errors.

To ensure the theoretical plausibility, parameters *A* and *B* were constrained between 0 and 1, where 1 represents perfect retention or full adaptation, and 0 represents complete decay or no adaptation, respectively. In contrast, parameter *C* was determined dynamically for each participant and each phase. The lower bound for *C* was set to the participant’s minimum observed reach angle minus a 5° margin, and the upper bound was set to their maximum observed reach angle plus a 5° margin. This dynamic approach ensured that the search space for parameter *C* was centered on each participant’s actual behavior, preventing the optimizer from settling on values far outside the observed performance range.

These models were selected based on the lowest Akaike Information Criterion (AIC) at the population level (Table S1). The goodness-of-fit and predictive accuracy for the winning models were also quantified using the R2 value and the Mean Squared Error (MSE; Table S1).

#### Model-free analysis

To complement the state-space model analysis and provide converging, model-free evidence, we quantified the speed of behavioral change during both adaptation and retention phases using two analogous metrics: Time to Adaptation and Time to Decay. For each participant and contingency, a participant-specific threshold was defined as 80% of their final performance level, computed as the average reach angle of the last twelve cycles of the respective phase. The metric was then calculated as the first cycle in which the participant’s reach angle crossed this threshold: exceeding it for adaptation or falling below it for retention. Lower values indicate faster adaptation or faster decay, respectively, reflecting more rapid learning or poorer retention.

#### Statistical analysis

To investigate the influence of reinforcement contingency on adaptation rate and retention, we compared the estimated parameters from the best-fitting state-space models across contingencies (Table S1). Specifically, we focused on the adaptation rate (B) from the AB model during the adaptation phase, and on the retention (A) and bias (C) parameters from the AC model during the retention phase. To rule out potential confounds, we also compared models’ fit quality and predictive accuracy (R² and MSE) across contingencies. In addition, we examined whether contingency effects changed depending on session order by splitting the data according to whether each contingency was experienced in the first or second session. All statistical tests between contingencies were conducted using paired t-tests or Wilcoxon signed-rank tests when normality assumptions were violated (Shapiro–Wilk test), with α = 0.05. The same analysis was used to assess the influence of feedback contingency on the model-free metrics (Time to Adaptation and Time to Decay) which were analyzed separately.

### Resting-state fMRI and behavior

We tested whether cerebello-cortical networks and neurotransmitter-enriched functional connectivity were associated with individual differences in adaptation rate under punishment and reward (Hypothesis 3A). Analyses were conducted separately for SERT and DAT, reflecting evidence that these neuromodulatory systems differentially influence reinforcement learning under reward and punishment [Cools et al., 2008; Cools et al., 2009; DenOuden et al., 2013; Doira et al., 2004; Palminteri et al., 2009; Palminteri et al., 2012; Worbe et al., 2011], which in turn shape adaptation rate [Galea et al., 2015]. For each neurotransmitter, connectivity values for CB6 and CBcrus1 served as dependent variables, and adaptation rates under punishment and reward were included as covariates. To capture whether the relationship between functional connectivity and adaptation differed across lobules depending on feedback type, a *Lobule* × *Contingency* interaction was included. Voxel-wise analyses were performed across the networks of interest (motor adaptation, reinforcement learning) using SPM12. Two complementary models captured alternative mappings between lobules and feedback conditions: (1) punishment primarily associated with CB6 and reward with CBcrus1, and (2) reward primarily associated with CB6 and punishment with CBcrus1. Statistical significance was set at *p* < 0.05 with FWE correction.

We finally tested whether the overlapping SERT and DAT network contributed to the retention of adapted movements (Hypothesis 3B), consistent with evidence that retention involves integrated neuromodulatory processes [Cardozo Pinto et al., 2025]. Multivariate linear regression (MLR) models were fitted separately for each ROI identified in the conjunction analysis (identifying overlapping SERT and DAT networks), with retention as the dependent variable and SERT- and DAT-enriched functional connectivity values, and their interaction as predictors (R; v4.3.2; R Development Core Team, 2023). All models included age, gender, and mean Framewise Displacement as covariates. The SERT × DAT interaction was tested using a likelihood ratio test (LRT), comparing models with and without the interaction term. When significant, post hoc pairwise comparisons of the association between DAT-connectivity values and retention (slopes β from the MLR) were performed at SERT-connectivity values corresponding to the mean, −1 SD, and +1 SD using the *emtrends* function in the *emmeans* package (v1.8.9), with Tukey’s method used to correct for multiple comparisons. Slopes (β) were reported along with their 95 percent confidence intervals (95% CI) and p-values. For interpretability, DAT and SERT measures were mean-centered and scaled to unit variance prior to analysis. All tests were two-sided, and statistical significance was set at p or adjusted *p* < 0.05.

## RESULTS

### Functional cerebello-cortical networks for CB6 and CBcrus1 have different spatial distributions

We examined whether SERT- and DAT-enriched networks were spatially distinct across cerebellar lobules (Hypothesis 1). Consistent with our prediction, the spatial distribution of DAT- and SERT-enriched networks differed between CB6 and CBcrus1. In CB6, DAT-enriched functional connectivity was higher than SERT-enriched functional connectivity within a selective set of nodes in the motor adaptation network (Figure 1A, *p* < 0.05, FWE correction). These nodes included the bilateral SMA, the dorsal premotor cortex, the hand representation of the primary motor cortex (M1) and the primary somatosensory cortex (S1), and the superior parietal lobule (Table S2). In contrast, SERT-enriched functional connectivity was higher with the ventral part of the motor adaptation network (Figure 1A), which included the inferior parietal lobule, the parietal operculum, and ventral premotor cortex (Table S2). No significant differences between SERT- and DAT-enriched functional connectivity were observed for the reinforcement network in CB6.

**Figure 1.**
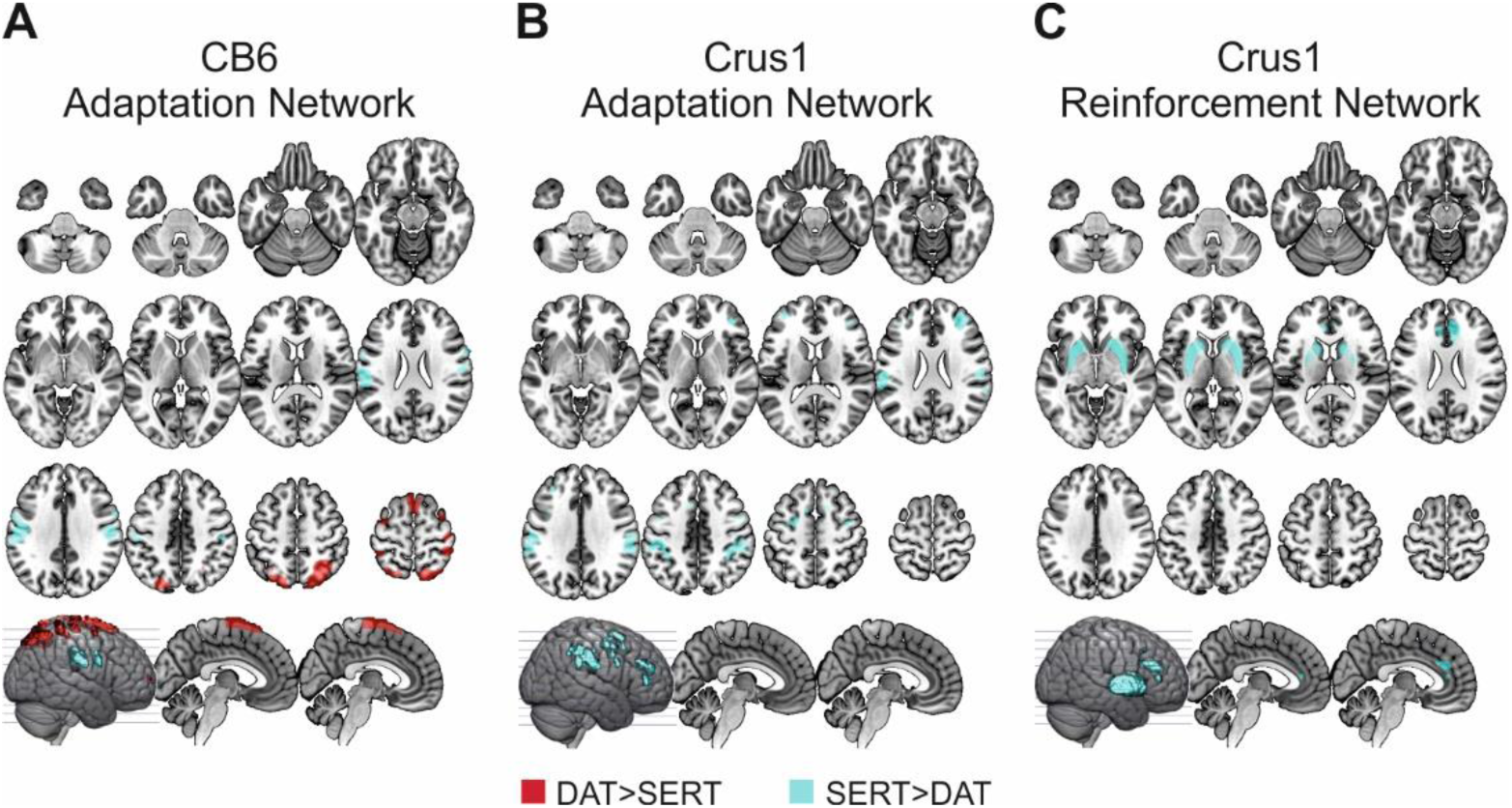
SERT- and DAT-enriched functional connectivity of cerebellar networks (*p* < 0.05, FWE correction). Group-level spatial covariance maps showing areas with increased functional connectivity with cerebellar lobules for DAT (red) or SERT (cyan). Group-level maps are overlaid on the anatomical MRI template in MRIcronGL: axial sections are shown from z = −45 to +65 mm in 10 mm increments for the first three rows, and sagittal sections are shown at x = −5, and x = +5 mm for the last row, with slices displayed on a 3D render. **A.** CB6 functional connectivity with the motor adaptation network. **B.** CBcrus1 functional connectivity with motor adaptation network. **C.** CBcrus1 FC with the reinforcement network.

In CBcrus1, SERT-enriched functional connectivity was higher than DAT-enriched functional connectivity within selective parts of both the motor adaptation and the reinforcement network (Figure 1B-C). For the motor adaptation network, these regions included the bilateral supplementary motor area, dorsal premotor cortex, hand representations of M1 and S1, and the inferior parietal lobule (Table S2). For the reinforcement network, SERT-enriched functional connectivity was stronger with the bilateral striatum (global maxima in ventral striatum corresponding to the nucleus accumbens) and bilateral medial prefrontal cortex, including anterior cingulate (BA 8 and 11; Table S2). No regions of the reinforcement network showed greater DAT-enriched functional connectivity compared with SERT-enriched functional connectivity in CBcrus1.

### Overlapping representation of SERT and DAT cerebello-cortical networks

We further tested whether resting-state SERT and DAT cerebello-cortical systems are functionally integrated within overlapping networks (Hypothesis 2). To this end, we performed conjunction analyses of SERT- and DAT-enriched connectivity maps for CB6 and CBcrus1, separately. As predicted, these analyses revealed regions in which cerebellar functional connectivity was jointly modulated by both neurotransmitters within each lobule. In CB6, the overlap was observed primarily within regions of the motor adaptation network, including the sensorimotor cortices and extended into the posterior parietal cortex (posterior part of the medial intraparietal sulcus integrating visual information and related to reaching movement; [Medendorp et al., 2005; Yttri et al., 2013]; Figure 2A, Table S3). In CBcrus1, the overlap was identified mainly within regions of the reinforcement network, including the bilateral NAC and medial orbitofrontal cortex (MOFC, Figure 2B, Table S3).

**Figure 2.**
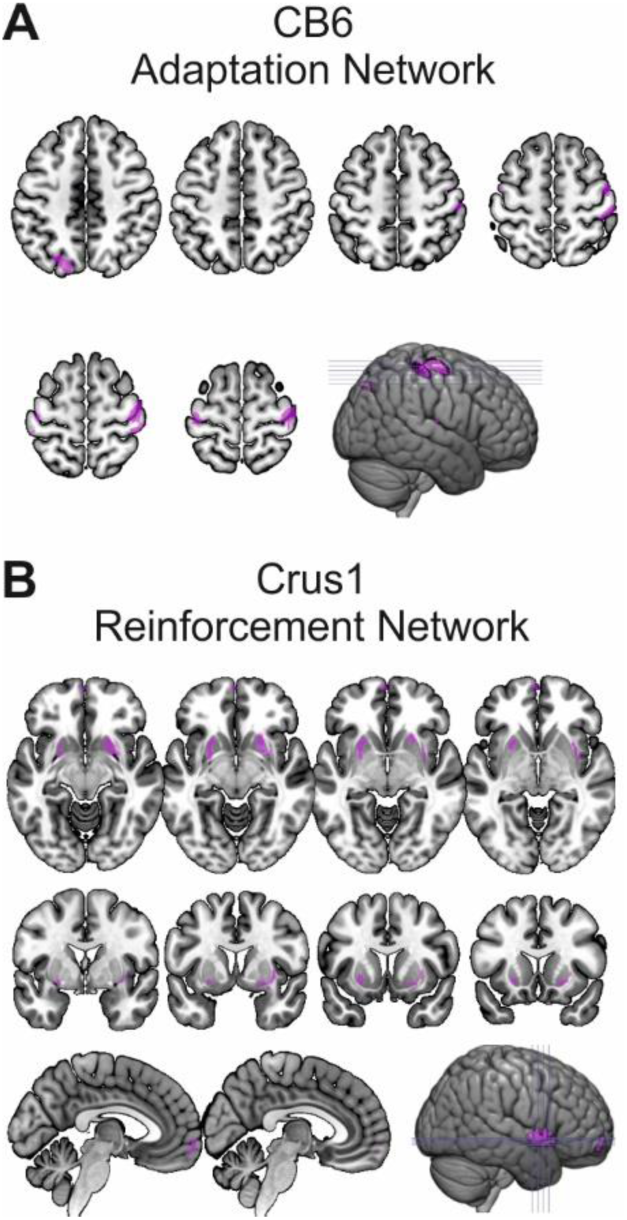
Result of the conjunction analysis showing the functional overlap between SERT- and DAT-enriched functional connectivity (*p* < 0.05, FWE correction). Group-level maps are overlaid on the anatomical MRI template in MRIcronGL. **A.** CB6 functional connectivity overlap within the motor adaptation network, shown on axial sections from z = +35 to +65 mm in 5 mm increments, with slices displayed on a 3D render. **B.** CBcrus1 functional connectivity overlap within the reinforcement network, shown on axial sections from z = −5 to +15 mm in 5 mm increments (first row) and coronal sections from y = 0 to +15 mm in 5 mm increments (second row), along with sagittal views from a 3D render and slices at z = −5 and +5 mm (last row).

### Punishment enhances adaptation rate compared to reward but retention is unaffected

We assessed the behavioral impact of reinforcement feedback on motor adaptation rate and retention. We observed a significantly higher adaptation rate under punishment compared to reward (*B* parameter: *W* = 246, *n* = 24, *z* = 2.74, *p* = 0.005, *rrb* = 0.64; Figure 3; also Figures S1 and S2 for individual adaptation profiles). In contrast, no significant differences emerged between punishment and reward for the retention parameters (*A* parameter: *W* = 164, *n* = 24, *z* = 0.40, *p* = 0.71, *rrb* = 0.09; *C* parameter: *W* = 147, *n* = 24, *z* = −0.86, *p* = 0.94, *rrb* = 0.02; Figure 3; also Figures S1 and S2 for individual retention profiles). This suggests that while punishment enhanced the speed of adaptation, it did not influence the retention or residual bias of the learned motor state. Importantly, these effects could not be attributed to differences in model fit quality, as neither MSE nor R2 differed across contingencies (all *p* > 0.06). Additionally, this effect was independent from the session order, as no significant differences were observed when comparing first- versus second-session exposures (all *p* > 0.12). Consistent with these model-based findings, the model-free analyses provided converging evidence: punishment led to faster adaptation (Time to Adaptation: *W* = 25, *n* = 24, *z* = −2.23, *p* = 0.02, *rrb* = −0.63), whereas retention was unaffected by contingency (Time to Decay: *W* = 174, *n* = 24, *z* = 0.70, *p* = 0.49, *rrb* = 0.16).

**Figure 3.**
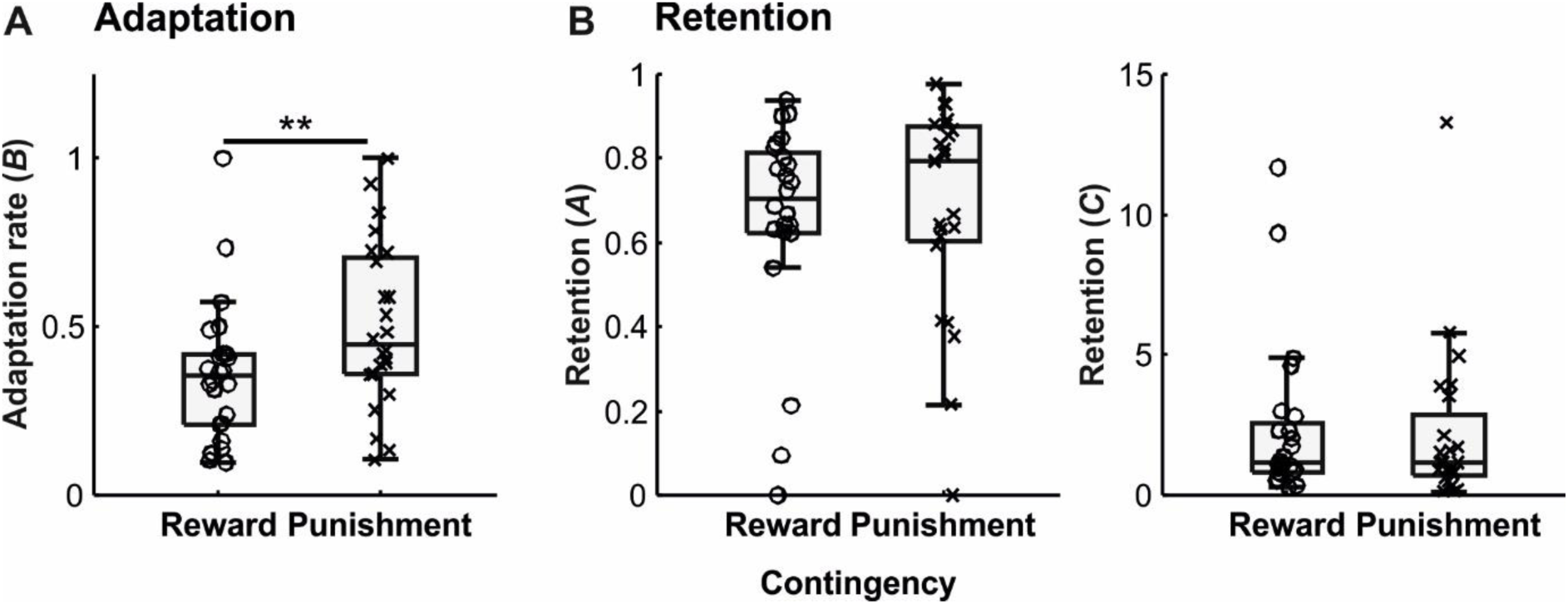
Distribution of participant values for **A.** Adaptation rate (*B)*, and **B.** Retention (*A* and *C*) for feedback contingencies (Reward *vs.* Punishment). Box charts represent the interquartile range (IQR; 25th to 75th percentiles) with the median indicated by the horizontal line. Whiskers extend to the most extreme data points within ±1.5 × IQR. Individual participant data are overlaid to show dispersion; dots represent the Reward contingency, while crosses represent the Punishment contingency. ** signifies *p* < 0.01. Individual profiles are shown in figures S1 (Reward) and S2 (Punishment).

### Distinct SERT- and DAT-enriched cerebello-cortical networks predict adaptation rates driven by reward and punishment

Having established the spatial organization and overlap of SERT- and DAT-enriched cerebello-cortical networks, we next examined whether these networks were related to individual differences in behavioral performance. Specifically, we tested whether neurotransmitter-enriched cerebello-cortical functional connectivity predicted adaptation rate under reward and punishment (Hypothesis 3A). First, DAT-enriched CB6-cortical connectivity was unrelated to adaptation rate, whereas DAT-enriched CBcrus1-SMA coupling was associated with punishment-driven adaptation rate (Figure 4A, Table S4). Specifically, greater DAT-enriched coupling between CBcrus1 and the supplementary motor area was associated with slower adaptation under punishment. Second, SERT-enriched cerebello-orbitofrontal coupling showed a positive correlation with adaptation rate under reward and punishment in CB6 and CBcrus1, respectively (Figure 4B, Table S4). Specifically, participants showing stronger SERT-enriched cerebello–orbitofrontal coupling adapted more rapidly in response to reward for CB6 but punishment for CBcrus1.

**Figure 4.**
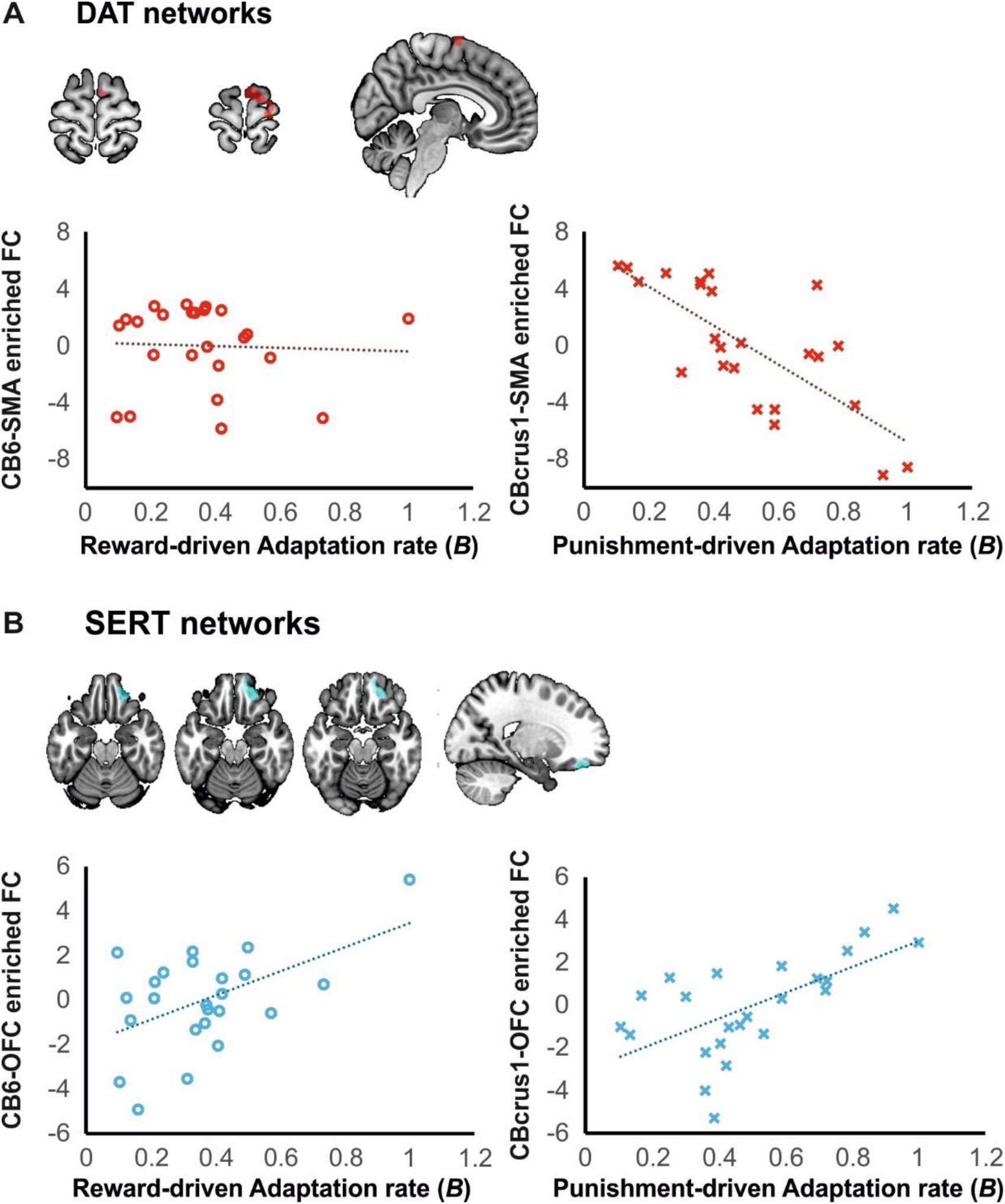
Relationship between DAT- and SERT-functional connectivity adaptation rates across feedback contingencies. Scatter plots illustrate the correlation between resting-state DAT- and SERT-enriched functional connectivity (FC) and individual adaptation rates (*B*) for Reward-driven and Punishment-driven adaptation. **A.** DAT networks: Correlation between cerebellum (CB6/CBcrus1) to SMA connectivity and adaptation. (Left) Reward-driven adaptation rate (represented by red circles) shows no significant correlation with CB6-SMA connectivity. (Right) Punishment-driven adaptation rate (represented by red crosses) shows a significant negative correlation with CBcrus1-SMA connectivity. Brain maps above indicate the ROI corresponding to the DAT network. **B.** SERT networks: Correlation between cerebellum (CB6/CBcrus1) to OFC connectivity and adaptation. (Left) Reward-driven adaptation rate (represented by blue circles) correlates positively with CB6-OFC connectivity. (Right) Punishment-driven adaptation rate (represented by blue crosses) correlates positively with CBcrus1-OFC connectivity. Brain maps above indicate the ROI corresponding to the DAT network. Dotted lines represent the linear regression fit. All behavioral data points (circles for Reward, crosses for Punishment) represent individual participant values (n = 24). Group-level maps are overlaid on the anatomical MRI template in MRIcronGL with the relevant clusters highlighted in red (DAT) and cyan (SERT).

### Interactions between SERT and DAT for CBcrus1 network explain inter-individual variability of retention behavior

We finally examined whether the interaction between SERT- and DAT-enriched networks could explain inter-individual variability in retention behavior (Hypothesis 3B). Because retention reflects longer-lasting processes that rely on integrated neuromodulatory mechanisms [Cardozo Pinto et al., 2025], we specifically focused on joint DAT–SERT effects. Interactions between DAT- and SERT-enriched functional connectivity in CBcrus1 predicted individual differences in retention following punishment and reward contingencies. In left PPC, the interaction was detectable only under the reward condition, where the LRT indicated a modest DAT × SERT effect (*p* = 0.031; Figure 5A) that did not survive pairwise correction in post hoc tests (0.08 < p < 0.83). In the MOFC, by contrast, a robust interaction emerged specifically under the punishment condition (LRT, *p* < 0.001; Figure 5B), reflecting a strong negative DAT-enriched connectivity slope at high SERT-enriched connectivity (+1 SD; *β* = −0.22, 95% CI [−0.32, −0.12], *p* < 0.001), a negative but non-significant slope at mean SERT levels (*β* = −0.06, 95% CI [−0.14, 0.01], *p* = 0.103), and a marginally positive trend at low SERT-enriched connectivity (−1 SD; *β* = 0.09, 95% CI [−0.01, 0.20], *p* = 0.084). All pairwise slope contrasts were significant (*p* < 0.001). Taken together, these findings indicate a clear punishment-specific SERT–DAT modulation in CBcrus1-MOFC coupling and a much subtler reward-related effect in left PPC. All remaining ROIs part of the conjunction analyses, including SMA, right striatum, and all other tested regions, showed no significant DAT × SERT interactions under either feedback contingency (all *p* > 0.051).

**Figure 5.**
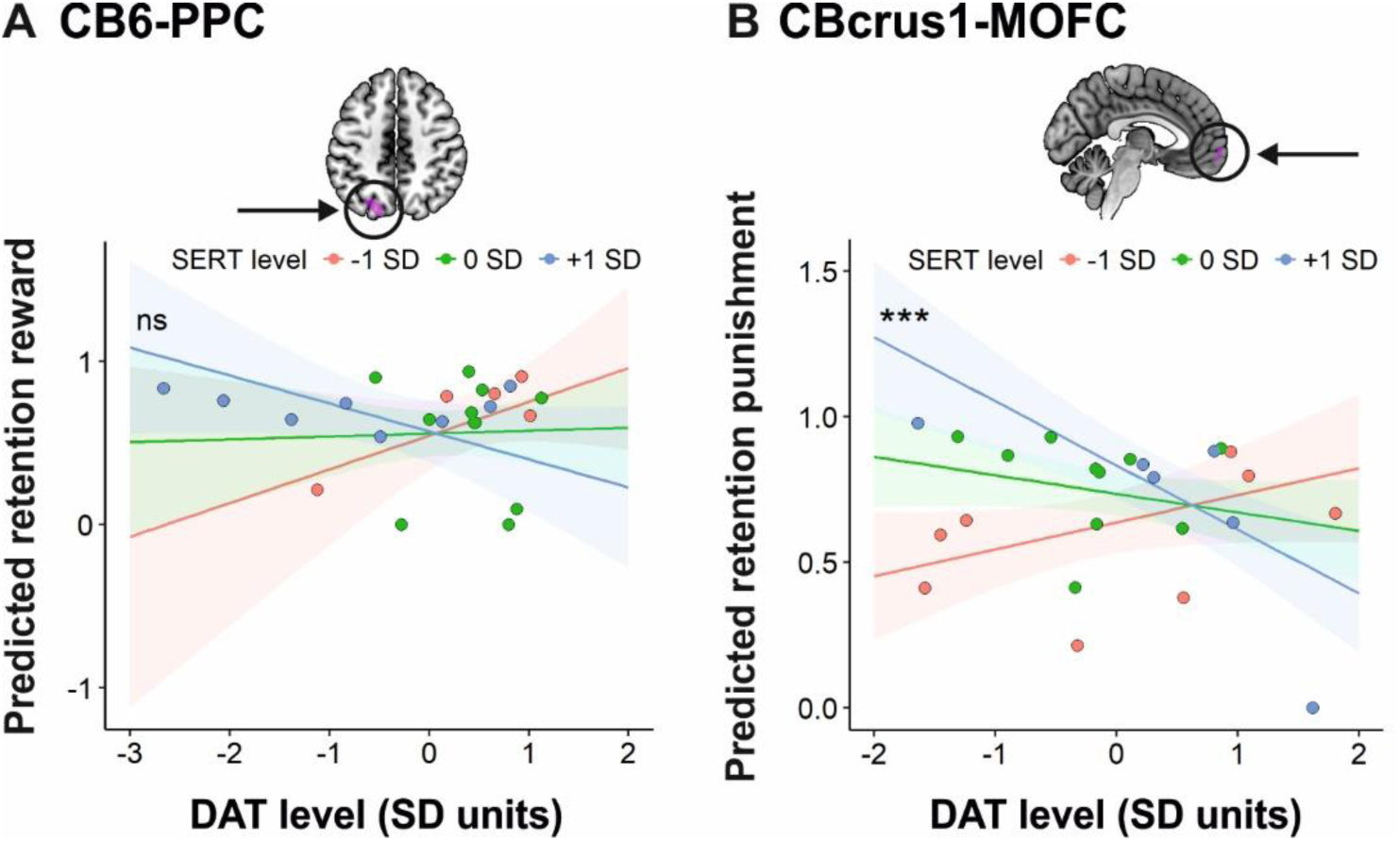
Relationship between DAT/SERT levels and predicted retention following reward and punishment. **A.** This panel illustrates the predicted retention following adaptation with reward, based on DAT levels across varying SERT levels within the CB6-PPC network as identified in the conjunction analysis (Figure2A). **B.** This panel illustrates the predicted retention following adaptation with punishment, based on DAT levels across varying SERT levels within the CBcrus1-MOFC network, as identified in the conjunction analysis (Figure2B). Predicted retention values are plotted against DAT levels (standard deviation units), with shaded areas representing the 95% confidence intervals for each SERT level group. Only results showing a significant DAT × SERT interaction are displayed. Ns (not significant) indicates that the post-hoc slopes did not differ statistically, whereas *** denotes a significant difference between all slopes with *p* < 0.001. Group-level conjunction maps are overlaid on the anatomical MRI template in MRIcronGL with the relevant clusters highlighted in pink.

## DISCUSSION

We investigated the distinct and joint contributions of serotonin and dopamine systems to reinforcement-based motor adaptation, focusing on the functional networks of CB6 and CBcrus1. We combined behavioral measures with SERT- and DAT-enriched resting-state functional connectivity using a template-based approach (REACT; [Dipasquale et al., 2019]). Our results provide converging evidence for functionally segregated and partially integrated neuromodulatory systems within cerebellar-cortical circuits, providing four main findings. First, we observed a marked differentiation between CB6 and CBcrus1: CB6 preferentially communicated with the motor adaptation network when functional connectivity was enriched both with SERT and DAT, whereas CBcrus1 communicated with both motor adaptation and reinforcement networks, especially when functional connectivity was SERT-enriched. Second, SERT- and DAT-enriched networks had overlaps expressed in adaptation regions for CB6 and in reinforcement processing regions for CBcrus1. Third, connectivity within CB6- and CBcrus1-enriched pathways explained the difference in adaptation performance depending on punishment- and reward-based adaptation. Finally, the interaction between the two neurotransmitter-enriched systems within the CBcrus1-prefrontal network predicted the retention of motor adaptation, particularly following punishment. Altogether, these findings indicate that serotonergic and dopaminergic cerebello–cortical networks are associated with reinforcement-based adaptive behavior at the interface of motor control and reinforcement processing.

### Topographical organization of neuromodulatory cerebellar networks

DAT- and SERT-enriched resting-state functional connectivity showed distinct spatial distributions of brain networks across cerebellar lobules. In CB6, DAT-enriched connectivity was strongest with dorsal cortical regions within the motor adaptation network, whereas SERT-enriched connectivity was strongest with ventral sensorimotor cortical regions. In contrast, CBcrus1 exhibited predominantly SERT-enriched connectivity spanning ventral sensorimotor cortical regions as well as those belonging to the reinforcement network. Despite this segregation, conjunction analyses revealed overlapping SERT- and DAT-enriched pathways in some cortical and subcortical nodes. Importantly, the spatial distribution of this overlap differed across cerebellar lobules: in CB6, it was observed primarily within regions of the motor adaptation network, while in CBcrus1, it was identified mainly within regions of the reinforcement network.

Together, these results suggest a topographical cerebellar organization in which CB6 is preferentially embedded within the sensorimotor cortical network, whereas CBcrus1 is more strongly coupled with associative and reinforcement-related cortical networks. These findings align with prior evidence for a rostro-caudal distribution in cerebellar functional organization, whereby CB6 is predominantly coupled to motor and premotor cortices, and CBcrus1 interfaces more with prefrontal and associative networks [Buckner et al., 2011; Diedrichsen et al., 2019; D’Mello et al., 2020; Nettekoven et al., 2024]. Within this framework, we bring additional information about possible factors contributing to this distribution, namely DAT and SERT neuromodulation. The differential contribution of DAT-enriched connectivity to dorsal motor networks and SERT-enriched connectivity to ventral sensorimotor and reinforcement-related networks is consistent with broader evidence that dopamine and serotonin differentially modulate large-scale resting-state networks by shaping subcortical-cortical functional connectivity. Specifically, dopaminergic systems are frequently linked to action-oriented/salience-related processing and sensorimotor network engagement, whereas serotonergic systems are often implicated in behavioral control and value- or context-dependent regulation within associative and reinforcement-related circuitry [Conio et al., 2020; Saiz-Masvidal et al., 2025]. Importantly, our findings further suggest that cerebellar-cortical circuits rely on the interaction of different neuromodulatory systems [Cardozo Pinto et al., 2025; González-Burgos and Feria-Velasco, 2008; Saiz-Masvidal et al., 2025; Sasaki-Adams and Kelley, 2001]. This segregation-integration pattern conceptually aligns with observations in Parkinson’s disease (PD), where both dopaminergic and serotonergic dysfunction contribute to symptom expression, with dopamine levels more closely linked to motor symptoms and serotonergic levels more strongly correlated with non-motor symptoms [Li et al., 2025]. From this perspective, the prominence of DAT-enriched connectivity in CB6-centered motor adaptation pathways, alongside stronger SERT-enriched connectivity in CBcrus1-centered associative/reinforcement pathways, is consistent with a partial motor–non-motor dissociation across dopamine and serotonin systems while still allowing convergence within shared circuits.

### Transporter-specific cerebellar pathways predict adaptation rate

The differential contribution of DAT- and SERT-enriched cerebellar pathways to motor adaptation and reinforcement networks was further supported by our brain-behavior results. At the behavioral level, we found that punishment led to faster adaptation than reward, consistent with prior work showing that negative feedback can accelerate motor adaptation [Galea et al., 2015]. At the brain level, we observed lobule- and neurotransmitter-specific associations with adaptation rate.

In CB6, faster adaptation under reward was associated with stronger SERT-enriched connectivity with orbitofrontal regions. This finding may be consistent with evidence that, in addition to its well-established sensorimotor functions [Buckner et al., 2011; Diedrichsen et al., 2019; Hardwick et al., 2013; Johnson et al., 2019], CB6 also contributes to reward-related processing [Chen et al., 2022; Kostadinov and Häusser, 2022; Wagner et al., 2017]. However, because reward did not confer a clear advantage to the adaptation rate, the observed CB6–orbitofrontal association may not reflect reward-driven adaptation per se. Instead, it may relate more generally to performance monitoring in motor adaptation. In this view, orbitofrontal involvement, often linked to outcome evaluation and flexible behavioral adjustment [Schoenbaum et al., 2009], suggests that serotonergic modulation of CB6 may support the cognitive aspects of motor adaptation, rather than its purely motor implementation [Boileau et al., 2008; Li et al., 2025].

In CBcrus1, punishment-driven adaptation showed dissociable associations across transporters and cortical pathways: stronger DAT-enriched connectivity with SMA was associated with slower adaptation, whereas stronger SERT-enriched connectivity with orbitofrontal regions was associated with faster adaptation. These opposing effects are consistent with evidence that dopamine-and serotonin-related systems can differentially shape learning and control processes, particularly under appetitive versus aversive contingencies [Colwell et al., 2024; Conio et al., 2020; González-Burgos and Feria-Velasco, 2008; Hamid et al., 2016; Miyazaki et al., 2014; Muehlberg et al., 2024; Sasaki-Adams and Kelley, 2001; Da Silva et al., 2018; Soubrié, 1986]. At the same time, the directionality of these associations should be interpreted with caution. Both neuromodulatory systems can exert facilitatory or suppressive influences depending on which receptors are engaged, their pre-/postsynaptic location, and the local microcircuit in the target region; moreover, serotonergic and dopaminergic signaling can reciprocally regulate each other’s release and postsynaptic impact [Abg Abd Wahab et al., 2019; González-Burgos and Feria-Velasco, 2008; Sasaki-Adams and Kelley, 2001]. Our approach indexes the spatial correspondence between functional connectivity and PET-derived transporter distributions (DAT and SERT). Accordingly, it cannot resolve receptor subtype contributions (e.g., D1-like vs D2-like; 5-HT1/2/4/6/7) or local excitation-inhibition balance. Additionally, given the polysynaptic and state-dependent nature of cerebello-thalamo-cortical loops, stronger coupling could reflect either facilitation of updating or increased stabilization/constraint, depending on the context [Allen and Tsukahara, 1974]. Based on our network-level results, one possible interpretation is that stronger DAT-informed CBcrus1–SMA coupling reflects a motor-biased configuration that favors slower adaptive stabilization over rapid updating under punishment; stronger SERT-informed CBcrus1–orbitofrontal coupling reflects a comparatively non-motor (evaluative/affective–cognitive) route supporting strategy-related adjustments that facilitate adaptation [Boileau et al., 2008; Li et al., 2025; Madsen et al., 2010].

### Integrated serotonin–dopamine contributions predict retention

In contrast with previous results, we did not detect a significant effect of reinforcement feedback on retention (i.e., retention did not differ between reward and punishment; [Galea et al., 2015]). Nevertheless, retention exhibited substantial inter-individual variability that could be captured by integrated SERT- and DAT-enriched connectivity effects. We assumed that retention would be characterized by networks showing a convergence of SERT- and DAT- influences (overlapping network) within the cerebello–cortical circuits, with areas included in the motor adaptation network for CB6 and reinforcement network for CBcrus1. Consistent with this account, retention following punishment was most strongly predicted by the interaction between DAT- and SERT-enriched CBcrus1 connectivity within the medial prefrontal cortex. This result mirrors the behavioral and neural findings for the adaptation rate, where connectivity within CBcrus1-centered associative networks was more strongly associated with punishment than reward. From a physiological perspective, this pattern is consistent with the idea that stabilization of learned behavior may rely on coordinated neuromodulatory influences within associative/prefrontal circuits, rather than being driven by either transmitter system alone [Cardozo Pinto et al., 2025]. One plausible mechanism for such coordination is that serotonergic signaling can shape dopaminergic effects in a receptor-dependent manner, with different 5-HT receptor subtypes exerting facilitatory or suppressive influences on dopamine release and downstream motor output [González-Burgos and Feria-Velasco, 2008; Sasaki-Adams and Kelley, 2001]. Although our approach cannot resolve receptor-level processes, the present effects are consistent with a model in which retention reflects a joint tuning of serotonergic and dopaminergic control within prefrontal and associative cerebellar networks. In this context, the dissociable dopaminergic and serotonergic associations observed during adaptation may linger during retention and be reorganized depending on the need to de-adapt when facing environmental changes.

### Limitations

Some limitations should be considered when interpreting the present findings. First, SERT- and DAT-enriched connectivity provides indirect estimates based on molecular PET-derived templates (n≈210–300) rather than participant-specific measures of transporter density or direct indices of neurotransmitter signaling. Although individual receptor binding potential maps would be ideal, they are invasive, costly, and ethically challenging in healthy participants. Moreover, DAT expression is relatively sparser at the cortical and cerebellar level relative to SERT, DAT-template enrichment may have offered a more limited dynamic range for detecting dopamine-linked effects outside classically DAT-rich territories, potentially contributing to weaker DAT-related findings. Second, resting-state networks were measured independently of task performance, with the behavioral task conducted outside the scanner. Although this design minimizes head motion and allows extensive trial repetition, it yields an indirect link between connectivity and behavior in a modest sample. Third, in contrast with previous results [Galea et al., 2015], we observed no significant effect of reward on retention of adapted movements. This likely reflects differences in experimental design: we used smaller visuomotor rotations (25°–30°) to limit inter-session savings in a within-subject design, which may have reduced reward-driven retention effects. Finally, cerebello-basal ganglia contributions may be underrepresented here, either because of the ROIs examined or because these interactions are primarily task-evoked and/or linked to behavioral dimensions not captured by our primary metrics.

## Conclusion

Our findings provide converging evidence that cerebello–cortical networks are topographically organized and differentially modulated by dopamine and serotonin to support reinforcement-based motor adaptation. CB6 primarily interfaces with motor cortical circuits through both DAT- and SERT-enriched connectivity, consistent with a motor–cognitive gradient in cerebellar organization. In contrast, CBcrus1 interfaces with associative and reinforcement networks and shows dissociable dopamine- and serotonin-linked associations with punishment-driven adaptation, highlighting that reinforcement-based motor learning engages cerebellar contributions, including evaluative and cognitive-control processes [Bracco et al., 2025; Diedrichsen et al., 2019; D’Mello et al., 2020; Kostadinov and Häusser, 2022; Kruithof et al., 2023; Taylor and Ivry, 2014]. Importantly, cerebellar influences during adaptation, particularly CBcrus1, are expressed as convergent (interaction) effects during retention within shared prefrontal–associative circuits, supporting an integrated neuromodulatory mechanism for stabilizing adapted behavior. More broadly, by linking cerebellar–cortical organization to coordinated serotonin–dopamine influences, our findings offer a systems-level framework that may help interpret learning and persistence abnormalities across neurological and psychiatric disorders in which the cerebellum is implicated and serotonergic–dopaminergic balance is altered (e.g., Parkinson’s disease and schizophrenia). Overall, our results demonstrate that adaptive motor control relies on both the segregation and integration of serotonergic and dopaminergic influences across cerebello–cortical systems, linking neuromodulation, feedback processing, and the retention of adaptive behavior.

## ACKNOWLEDGMENTS

This work was carried out on the MRI and PANAM platforms of the neuroimaging center (Center de Neuroimagerie de Recherche; CENIR) within the Paris Brain Institute (ICM). This study was supported by the European Union’s Horizon 2020 research and innovation program under the Marie Skłodowska-Curie grant agreement 897941 to MB and the Big Brain Theory (BBT-3; FORTE) internal ICM grant to MB, AV-C and CG.

## AUTHOR CONTRIBUTIONS

Conceptualization: MB, CG. Methodology: MB, BB, AVC, CG. Software: SO, BB, KN. Investigation: MB, VR, XCT, AP, OO. Formal analysis: MB, CA, OO, FXL, SO, CG. Data Curation: MB. Writing—original draft: MB, CG. Writing—review & editing: MB, CA, VR, XCT, AP, OO, FXL, SO, BB, KN, YW, TP, AVC, CG. Visualization: MB, CG. Supervision: AVC, CG. Project administration: MB, AVC, CG. Funding acquisition: MB, AVC, CG.

