## Supplemental Figures and Tables for "Functional cerebellar connectomes interfacing motor adaptation and reinforcement feedback"

[*] Corresponding author

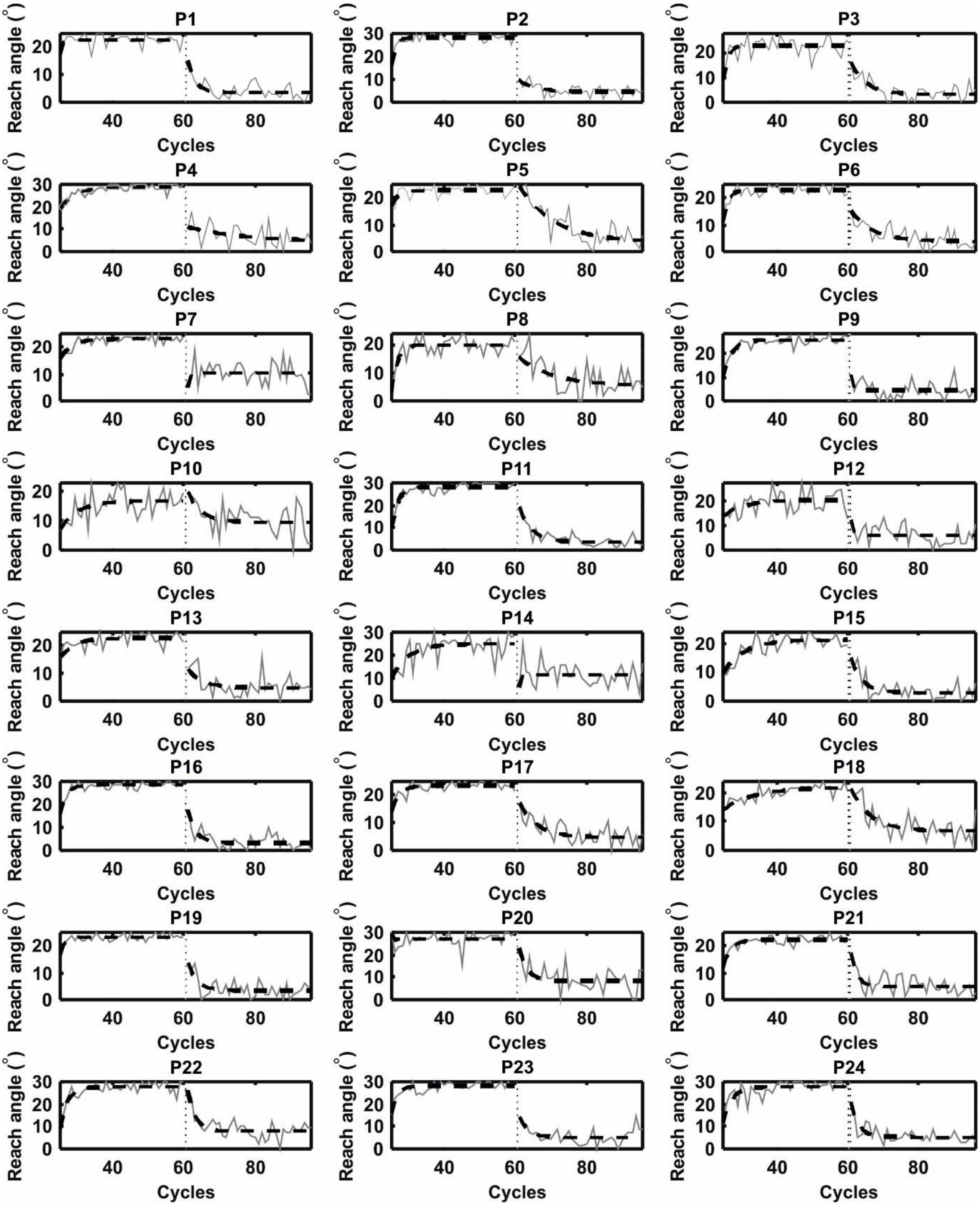


**Figure S1. Individual adaptation and retention profiles with State-Space Model (SSM) predictions during the Reward session.** Reach angle errors are plotted for each of the 24 participants across the adaptation and retention blocks. Data are presented in cycles, where each cycle represents the mean of four consecutive trials containing one reach to each of the four target positions. The continuous gray lines represent the observed (real) reach angle error, while the dashed black lines indicate the error predicted by the SSM fit. The transition from the adaptation to the retention block is marked by the vertical dashed line.

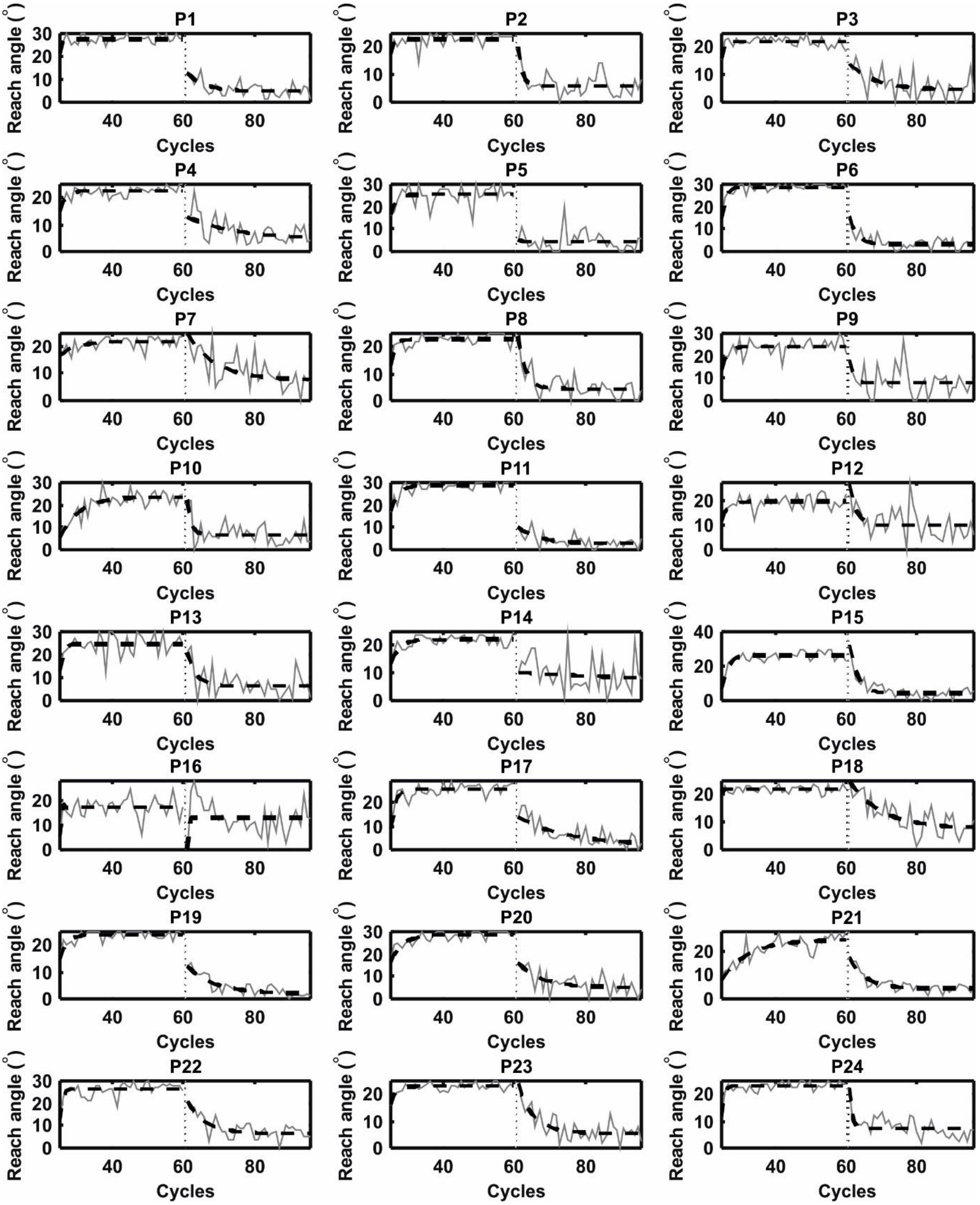


**Figure S2. Individual adaptation and retention profiles with State-Space Model (SSM) predictions during the Punishment session.** Reach angle errors are plotted for each of the 24 participants across the adaptation and retention blocks. Data are presented in cycles, where each cycle represents the mean of four consecutive trials containing one reach to each of the four target positions. The continuous gray lines represent the observed (real) reach angle error, while the dashed black lines indicate the error predicted by the SSM fit. The transition from the adaptation to the retention block is marked by the vertical dashed line.

**Table S1. Table S3. Model comparison for Reward and Punishment conditions across adaptation blocks (Adaptation *vs.* Retention)**. Values represent the mean ± standard deviation (SD) for each metric. (A) Goodness-of-fit was evaluated using Mean Squared Error (MSE), Coefficient of Determination ($R^2$), and Akaike Information Criterion (AIC). Two computational models were compared: the AB Model and the AC Model. Asterisks (*) denote the best-fitting model for each condition and adaptation block, identified by the lowest MSE and AIC values and the highest $R^2$.

| Condition | Phase | Model | MSE (± SD) | $R^2$ (± SD) | AIC (± SD) |
|---|---|---|---|---|---|
| **Reward** | Adaptation | AB Model* | 4.93 ± 4.50 | 0.548 ± 0.218 | 50.07 ± 27.84 |
| | | AC Model | 5.11 ± 4.54 | 0.523 ± 0.223 | 52.01 ± 27.09 |
| | Retention | AB Model | 17.67 ± 11.01 | −0.068 ± 0.531 | 102.40 ± 18.32 |
| | | AC Model* | 10.02 ± 4.93 | 0.403 ± 0.211 | 82.80 ± 17.83 |
| **Punish.** | Adaptation | AB Model* | 5.15 ± 4.04 | 0.453 ± 0.213 | 53.61 ± 26.10 |
| | | AC Model | 5.18 ± 4.02 | 0.447 ± 0.215 | 53.97 ± 26.02 |
| | Retention | AB Model | 27.48 ± 39.44 | −0.052 ± 0.975 | 108.88 ± 29.18 |
| | | AC Model* | 14.13 ± 9.44 | 0.422 ± 0.225 | 91.28 ± 25.56 |

**Table S2. Post-Hoc t-tests on functional connectivity maps of cerebellar lobules.** Statistics are shown for clusters with $p < 0.05$ with FWE correction on the volume of the network mask. Spatial representation of clusters is illustrated in Figure 1.

| **Anatomical localization of global maxima and cluster extent** | **Coordinates** | | | **Z-score (T when infi-nite)** | **Cluster volume** |
|---|---|---|---|---|---|
| | **x** | **y** | **z** | | |
| **CB6: DA > SERT (Adaptation Network)** | | | | | |
| Right posterior parietal cortex (superior parietal lobule, BA 5/7) | 6 | -68 | 66 | 12.36 | 610 |
| Right medial precentral gyrus (BA 6, SMA) | 4 | -15 | 66 | 11.27 | 701 |
| Left posterior parietal cortex (superior parietal lobule, BA 5/7) | -6 | -82 | 50 | 10.48 | 181 |
| Right postcentral gyrus (BA 1/2, somatosensory cortex) | 9 | -42 | 70 | 6.56 | 10 |
| Left lateral precentral gyrus (BA 6, PMd) | -36 | -5 | 66 | 6.19 | 55 |
| Left lateral postcentral gyrus (BA 1/2/3, soma-tosensory cortex) | -36 | -42 | 68 | 5.80 | 39 |
| Right lateral precentral gyrus (BA 6, PMd) | 37 | 0 | 66 | 5.21 | 32 |
| **CB6: SERT > DA (Adaptation Network)** | | | | | |
| Left inferior postcentral gyrus (BA 2/3) with ex-tension into inferior precentral gyrus (BA 6, PMv) | -54 | -20 | 33 | 5.58 | 558 |
| Right inferior postcentral gyrus (BA 2/3) | 56 | -20 | 31 | 4.85 | 219 |
| **CBcrus1: SERT > DA (Adaptation Network)** | | | | | |
| Left supramarginal gyrus (BA 40) | -58 | -32 | 28 | 6.48 | 456 |
| Right supramarginal gyrus (BA 40) | 59 | -22 | 33 | 5.96 | 384 |
| Right superior middle frontal gyrus (BA 6/8) | 36 | -8 | -50 | 5.82 | 54 |
| Right superior frontal gyrus (BA 8/6, FEF/SEF) | 16 | 10 | 60 | 5.63 | 67 |
| Right dorsolateral prefrontal cortex (BA 10/46) | 34 | 38 | 26 | 5.60 | 160 |
| Left middle frontal gyrus (BA 6/8) | -24 | -5 | 53 | 5.28 | 149 |
| Left dorsolateral prefrontal cortex (BA 10/46) | -36 | 48 | 13 | 5.26 | 118 |
| Right inferior frontal gyrus (BA 11/47) | 39 | 42 | 6 | 5.13 | 47 |

| | | | | | |
|---|---|---|---|---|---|
| Right inferior middle frontal gyrus | 54 | 2 | 40 | 5.13 | 26 |
| Left superior frontal gyrus (BA 8/6, FEF/SEF) | -8 | 18 | 43 | 4.93 | 32 |
| **CBcrus1: SERT > DA (Reinforcement Network)** | | | | | |
| Left putamen | -24 | 10 | 8 | 9.59 | 546 |
| Right putamen | 22 | 12 | 8 | 9.35 | 625 |
| Right anterior cingulate cortex | 12 | 28 | 28 | 6.48 | 177 |
| Left anterior cingulate cortex | -8 | 22 | 30 | 5.09 | 104 |
| Left dorsal anterior cingulate cortex | -8 | 18 | 40 | 5.01 | 8 |
| Right ventral anterior cingulate cortex | 9 | 38 | 8 | 5.54 | 21 |
| Right dorsal anterior cingulate cortex | 6 | 20 | 43 | 4.49 | 4 |

**Table S3. Conjunction analyses on functional connectivity maps of cerebellar lobules.** Statistics are shown for clusters with $p < 0.05$ with FWE correction on the volume of the network mask. Spatial representation of clusters is illustrated in Figure 2.

| **Anatomical localization** | **Coordinates** | | | **Z-score** | **Cluster volume** |
|---|---|---|---|---|---|
| | **x** | **y** | **z** | | |
| **CB6: DAT ∩ SERT (Motor Adaptation Network)** | | | | | |
| Right medial precentral gyrus (BA 6, SMA) | -6 | -8 | 75 | 5.88 | 112 |
| Right lateral precentral gyrus (BA 4/6, M1/PMd) | 42 | -18 | 63 | 5.78 | 181 |
| Right somatosensory cortex (BA 2) | 56 | -21 | 53 | 5.23 | 181 |
| Left posterior intraparietal sulcus (BA 7) | -23 | -72 | 43 | 5.76 | 24 |
| Left somatosensory cortex (BA 2) | -48 | -32 | 60 | 5.38 | 45 |
| Left lateral precentral gyrus (BA 4/6, M1/PMd) | -46 | -7 | 55 | 4.97 | 12 |
| **CBcrus1: DAT ∩ SERT (Reinforcement Network)** | | | | | |
| Right ventral putamen | 24 | 5 | -10 | 5.32 | 64 |
| Left ventral putamen | −24 | 0 | −9 | 5.07 | 39 |
| Medial orbitofrontal cortex (BA 11) | −1 | 60 | −14 | 4.49 | 24 |

**Table S4. Interaction between cerebellar functional connectivity and adaptive behavior.** Functional connectivity maps of cerebellar lobules explained the difference of adaptation rate under reward and punishment. Statistics are shown for clusters with $p < 0.05$ with FWE correction on the volume of the network mask. Spatial representation of clusters is illustrated in Figure 4.

| **Anatomical localization** | **Coordinates** | | | **F (Zscore)** | **Cluster volume** |
|---|---|---|---|---|---|
| | **x** | **y** | **z** | | |
| **DAT: $CB6_{reward}$ < $CBcrus1_{punishment}$ (Motor Adaptation Network)** | | | | | |
| Right medial precentral gyrus (BA 6, SMA) | 6 | -2 | 73 | 14.23 | 83 |
| **SERT: $CB6_{reward}$ < $CBcrus1_{punishment}$ (Reinforcement Network)** | | | | | |
| Right medial orbitofrontal cortex (BA 11, OFC) | 19 | 45 | -76 | 11.00 | 95 |
